# Specific ZNF274 binding interference at *SNORD116* activates the maternal transcripts in Prader-Willi syndrome neurons

**DOI:** 10.1101/2020.06.13.149864

**Authors:** Maéva Langouët, Dea Gorka, Clarisse Orniacki, Clémence M Dupont-Thibert, Michael S Chung, Heather R Glatt-Deeley, Noelle Germain, Leann J Crandall, Justin L Cotney, Christopher E Stoddard, Marc Lalande, Stormy J Chamberlain

**Affiliations:** Department of Genetics and Genome Sciences, School of Medicine, University of Connecticut, Farmington CT, 06013, USA; Institute for Systems Genomics, University of Connecticut, Farmington CT, 06013, USA

## Abstract

Prader-Willi syndrome (PWS) is characterized by neonatal hypotonia, developmental delay, and hyperphagia/obesity. This disorder is caused by the absence of paternally-expressed gene products from chromosome 15q11-q13. We previously demonstrated that knocking out ZNF274, a KRAB-domain zinc finger protein capable of recruiting epigenetic machinery to deposit the H3K9me3 repressive histone modification, can activate expression from the normally silent maternal allele of *SNORD116* in neurons derived from PWS iPSCs. However, ZNF274 has many other targets in the genome in addition to *SNORD116*. Depleting ZNF274 will surely affect the expression of other important genes and disrupt other pathways. Here we used CRISPR/Cas9 to delete ZNF274 binding sites at the *SNORD116* locus to determine whether activation of the maternal copy of *SNORD116* could be achieved without altering ZNF274 protein levels. We obtained similar activation of gene expression from the normally silenced maternal allele in neurons derived from PWS iPSCs, compared to ZNF274 knockout, demonstrating that ZNF274 is directly involved in the repression of *SNORD116*. These results suggest that interfering with ZNF274 binding at the maternal *SNORD116* locus is a potential therapeutic strategy for PWS.

## Introduction

Prader-Willi syndrome (PWS; OMIM 176270) is a neurogenetic disorder of genomic imprinting and has an incidence of ∼1/15,000 live births. Children affected with PWS suffer neonatal hypotonia and failure-to-thrive during infancy, followed by hyperphagia/obesity; small stature, hands, and feet; mild to moderate cognitive deficit; and behavioral problems that are likened to obsessive-compulsive disorder. PWS most commonly results from large deletions mediated by repetitive sequences flanking a ∼5 Mb imprinted region on paternal chromosome 15q11-q13^1; 2^. There is no cure for PWS. Current treatments focus on alleviation of individual symptoms^3-8^.

Many genes in the chromosome 15q11-q13 region are regulated by genomic imprinting. Most genes, including *SNRPN* (a bicistronic transcript that also encodes *SNURF*, referred to henceforth as *SNRPN* only), *SNHG14, MKRN3, MAGEL2*, and *NDN* are exclusively expressed from the paternally inherited allele. *UBE3A* is biallelic in most tissues, but in neurons, this gene is expressed from the maternally inherited allele only. *SNHG14*, a long non-coding RNA (*lncRNA*) initiated at the canonical and upstream promoters of *SNRPN* on the paternal allele (Fig. 1), extends >600kb distally and overlaps *UBE3A*, therefore silencing the paternal *UBE3A* allele^9-17^. *SNHG14* also serves as the host gene (HG) to several box C/D class small nucleolar RNAs, organized in large, tandemly repeated clusters, known as the *SNORD116* and *SNORD115* clusters^9; 17^. The 30 copies of the *SNORD116* cluster have been subdivided into 3 groups based on DNA sequence similarity^18^; Group 1 (*SNOG1, SNORD116 1-9)*, Group 2, (*SNOG2, SNORD116 10-24*) and Group 3 (*SNOG3, SNORD116 25-30)*. The PWS-Imprinting Center (PWS-IC), a region of differential CpG methylation, located in the promoter and first exon of *SNRPN*, is known to control imprinting at this region^19^.

**Figure 1.**
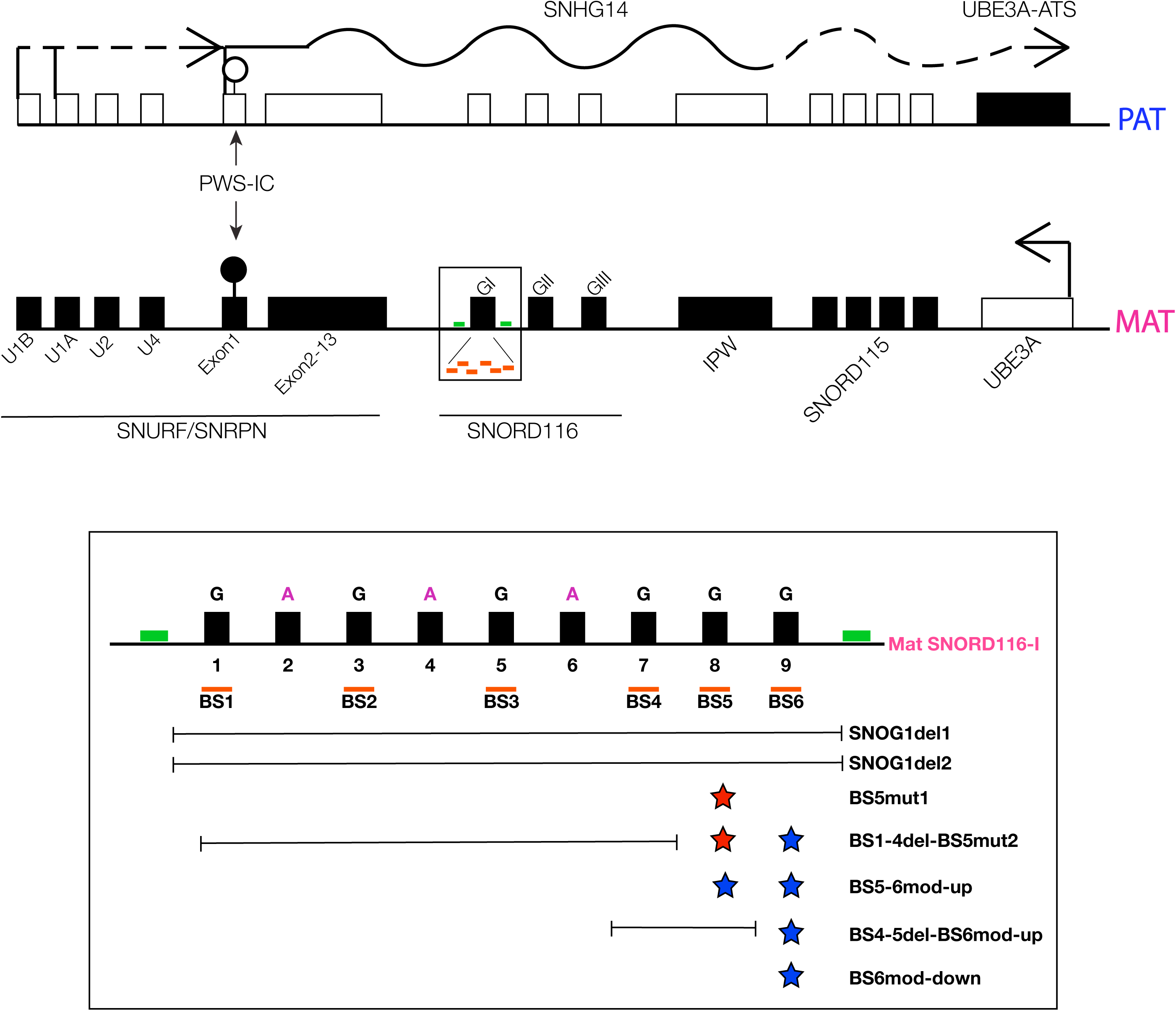
Summary of ZNF274 binding site modifications at the *SNORD116* locus. Simplified map of 15q11.2-q13. Active and inactive transcripts are denoted by open and closed boxes, respectively. Arrows indicate the direction of transcription. A solid black line represents paternal *SNHG14* transcript expressed in most cell types, whereas a dashed black line indicates neuron-specific transcripts, including upstream exons of *SNRPN* and *UBE3A-ATS*. The PWS-IC is denoted by the black (methylated)/white (un-methylated) circle. Orange dashes under the *SNORD116* cluster represent the six ZNF274 binding sites within the *SNORD116*s classified as Group 1 (*SNOG1-BS1* to *SNOG1-BS6*). Positions of SNOG1del Guide-1 and -2 are indicated with green dashes, surrounding *SNORD116*. In the zoomed area below, positions of large deletions spanning multiple or all the 6 ZNF274 Binding sites are indicated, as well as each mutation (red star) or modification (blue star) described in each cell line generated in this paper.

Although the genes involved in PWS have been known for many years, the exact contribution of each gene to the symptoms of PWS remain unclear. Efforts have been made to elucidate the targets of PWS snoRNAs: *SNORD115* is thought to regulate splicing of the serotonin HTR2C receptor ^20; 21^ and *SNORD116* has been computationally predicted to interact with *ANKRD11* mRNA, and perhaps other transcripts^20^. Additionally, Keshavarz et al demonstrated a correlation between copy number variation of *SNORD115* and *SNORD116* and behavioral traits, by assessing anxiety both in rodents and humans^22^.

In the past decade, focus has shifted to *SNORD116* because recently identified patients with atypical, shorter deletions suggest that most features of PWS could result from the loss of the *SNORD116* snoRNA cluster^23-26^. Additionally, mouse models produced by deletion of the *Snord116* cluster show several features of PWS including postnatal growth retardation, increased body weight gain and hyperphagia, further supporting the association between *Snord116* and PWS^27-29^. Moreover, recent work also demonstrated that loss of *SNORD116* in both human induced pluripotent stem cell (iPSC) and mouse models of PWS can lead to a deficiency of prohormone convertase PC1, an intriguing observation that may link *SNORD116* to the neuroendocrine dysfunction in PWS^30; 31^. However, whether the absence of *SNORD116* genomic region alone, its host-gene lncRNA transcript, the processed snoRNAs, and/or simply the active transcription event itself rather than the genomic region/RNA products is responsible of the disease remains an active debate.

Since every individual with PWS has a functional copy of the genetic region that is epigenetically silenced, activation of these genes offers an attractive therapeutic approach for this disorder. Using our PWS and Angelman Syndrome (AS) iPSC models, we previously reported that the KRAB-domain zinc finger protein ZNF274 binds to six sites on the maternal copy of the *SNORD116* cluster where it associated with the histone methyltransferase, SETDB1, and mediates the deposition of the repressive H3K9me3 chromatin mark on the maternal allele.^32-34^ By knocking out *ZNF274*, we were able to activate the silent maternal allele in PWS iPSC-derived neurons, without affecting DNA methylation at the PWS-IC.^35^ These results suggested that the ZNF274 complex mediates a separate imprinting mark that represses maternal PWS gene expression in neurons. Genome-wide *ZNF274* depletion, however, does not represent an ideal therapeutic strategy since ZNF274 may have crucial functions outside the PWS locus.^36^ Here we deleted and mutated the ZNF274 binding sites (BS) within the *SNORD116* locus in human PWS induced pluripotent stem cells (iPSCs). We found that preventing ZNF274 from binding leads to activation of maternal copies of PWS genes in human PWS iPSC-derived neurons. This demonstrates that *SNORD116* is a direct target of ZNF274-mediated repression. A strategy to inhibit binding of ZNF274 specifically at the maternal *SNORD116* region could potentially restore gene expression from the maternal copies of the PWS genes, while not affecting the other ZNF274-bound loci, providing what may be an optimal therapeutic approach for PWS.

## Results

### Identification of the *ZNF274* consensus binding motif

In order to design strategies to block ZNF274 binding at *SNORD116*, we developed a computational approach to search for a consensus DNA binding site for ZNF274. We analyzed 21 ZNF274 chromatin immunoprecipitation followed by sequencing (ChIP-Seq) datasets from 8 different cultured cell lines performed by the ENCODE Consortium and identified 1572 reproducibly bound sites in the human genome. We extracted the sequence of each of these sites from the reference human genome and analyzed this set with the Multiple Em for Motif Elicitation (MEME) suite^37^. We were able to identify a single binding motif for ZNF274 (Fig. 2A). Using this consensus binding motif, we then predicted all ZNF274 binding sites genome-wide using the Find Individual Motif Occurences (FIMO)^38^ routine from the MEME suite ^37^. The best match to the consensus ZNF274 motif elicited from ChIP-Seq data (TGAGTGAGAACTCATACC) was identified five times within the *SNORD116* cluster (Fig. 3A). Another group independently identified a putative ZNF274 binding motif.^39^ This motif is similar to ours, and is only shifted 2 bp downstream (Fig. 3A). The *SNORD116* cluster is comprised of 30 copies of the snoRNA and can be classified into 3 groups based on DNA sequence similarity^18^. Group 1 consists of *SNORD116-1* through *SNORD116-9* (Fig. 1). The exact ZNF274 motif was identified in five of the nine copies of *SNORD116* within this group, *SNORD116*-3,-5,-7,-8, and -9 (Fig. 2B). *SNORD116*-1 contains a single nucleotide change (at position 17) from the ZNF274 consensus binding motif (Fig. 3A). ChIP-Seq data indicates that the binding here is less reproducible, suggesting that this single nucleotide change may reduce ZNF274 binding affinity (Fig. 2B). Nonetheless, in human pluripotent stem cells, ZNF274 binds to all six predicted ZNF274 binding sites within *SNORD116*, as determined by ChIP-seq and ChIP-qPCR ^32; 35^, despite the single nucleotide change. *SNORD116*-2, -4, and -6 each display a G-to-A substitution at position 8 in the consensus motif (in magenta, Fig. 3A) and were not identified as being bound by ZNF274 in ChIP-Seq data. The consensus binding motif was also found in all nine Group 1 *SNORD116* copies in the cynomolgous monkey (*Macaca fascicularis*) genome, albeit without the A-to-G change. We confirmed ZNF274 binding at three *SNORD116* copies in cynomolgous iPSCs by ChIP-qPCR (Fig. 2C). This demonstrates the conservation of the ZNF274 consensus binding motif in primates.

**Figure 2.**
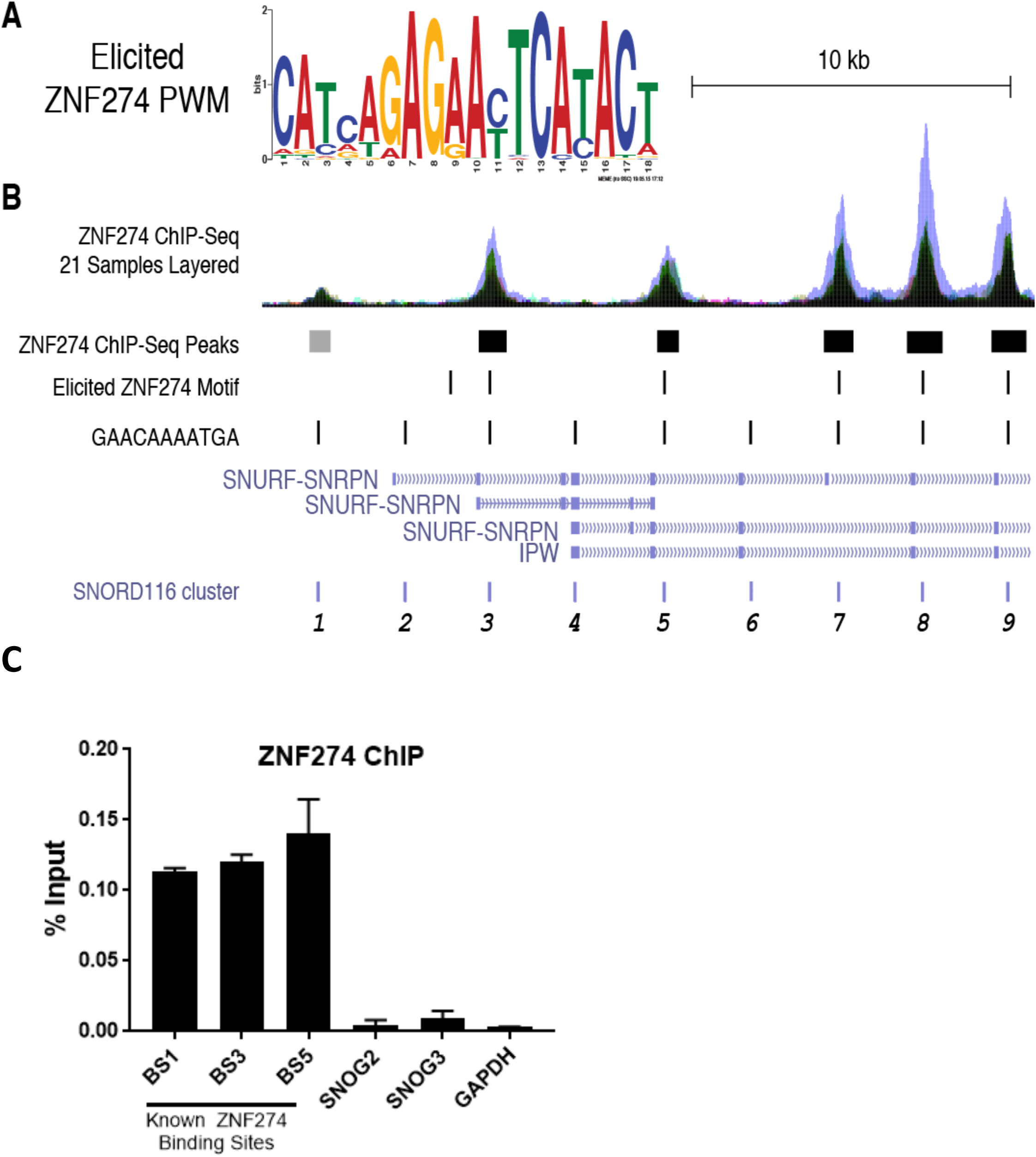
Region of nucleotide homology surrounding the ZNF274 motif at *SNORD116*. **A**. ZNF274 PWM elicited from over 1500 highly reproducible binding sites. **B**. ENCODE ZNF-274 ChIP-Seq composite signal and peak calls at *SNORD116-1,-3,-5,-7,-8,-9*. Boxes below signal tracks indicate binding sites. **C**. ZNF274 ChIP assays for cynomolgus stem cells.

**Figure 3.**
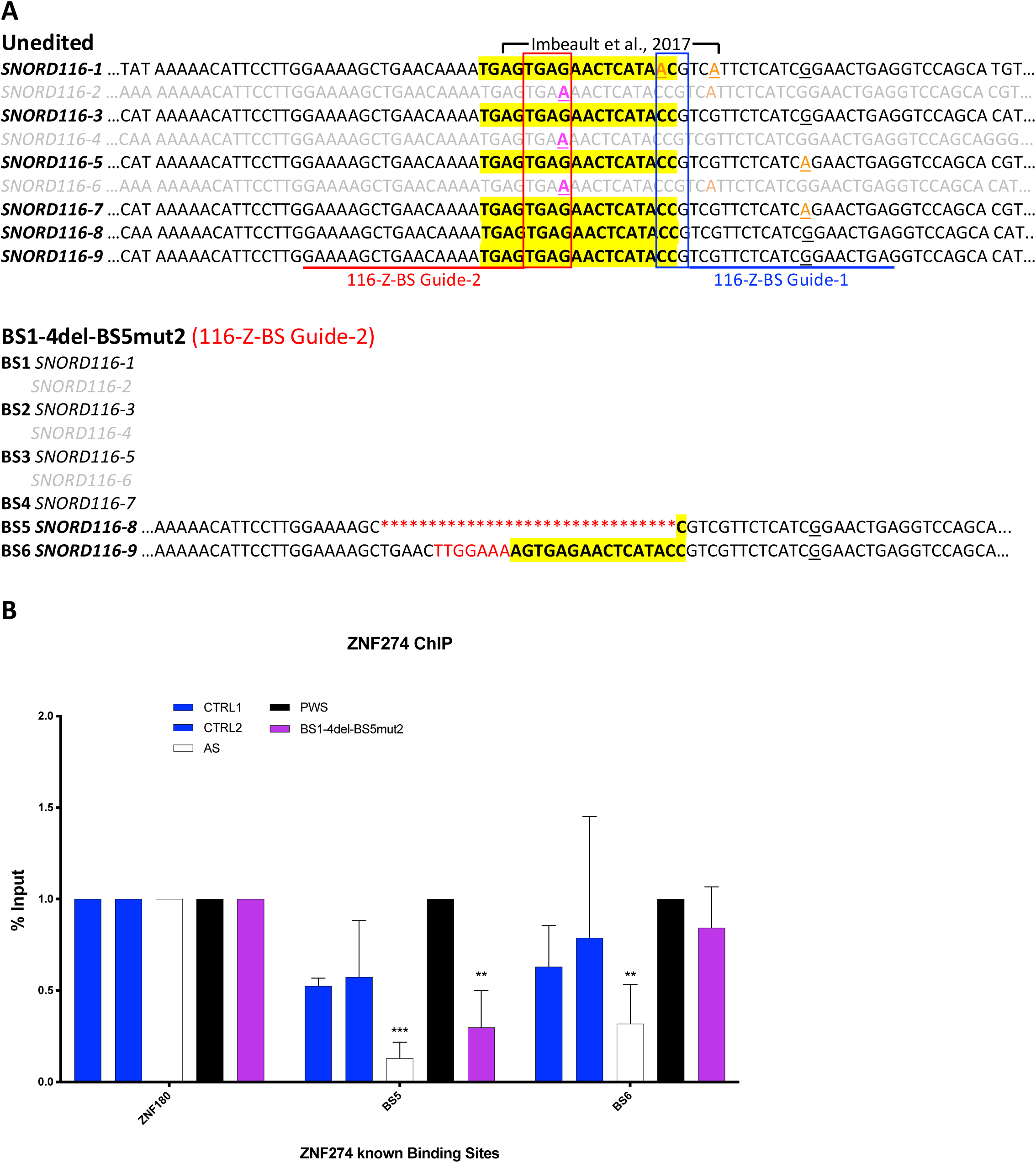
ZNF274 binding at *SNORD116*. **A**. DNA sequences of portions of group 1 *SNORD116-1* through *SNORD116-9* are shown. The ZNF274 consensus sequence identified herein is highlighted in yellow. The position of the ZNF274 motif proposed by Imbeault et al. is indicated. *SNORD116* copies bound by ZNF274 are in black font, while those not bound by ZNF274 are in gray font. Single base substitutions are highlighted in colored fonts. The positions of gRNAs targeting ZNF274 binding sites at *SNORD116* are underlined in blue and red. Their respective PAM sequences are in boxes. Lower panel illustrates the mutation sustained in one of the mutated clones to show the genetic alterations incurred at each ZNF274 binding site. **B**. ChIP-qPCR for ZNF274 in iPSCs. Quantification of ChIP was performed and calculated as percent input for each sample. Binding at *ZNF180* is included as a positive control. Samples were normalized against the PWS (black) sample. A minimum of 2 biological replicates per cell line were performed. Significance was calculated using two-way analysis of variance (ANOVA) test with a Dunnett post-test to compare the disrupted ZNF274 binding cell lines to PWS. *P<0.05, **P<0.01, ***P<0.001, ****P<0.0001.

### Generation of PWS iPSCs cell lines with modified ZNF274 binding sites

We sought to determine whether disruption of the ZNF274 binding sites within the *SNORD116* cluster would lead to activation of maternal *SNORD116* in neurons derived from PWS iPSCs. First, we used CRISPR/Cas9 to delete the entire cluster of six ZNF274 binding sites in PWS iPSCs harboring a large deletion of paternal 15q11-q13. We designed two guide RNAs (gRNAs) - SNOG1del Guide-1 is 5’ to binding site 1 (BS1) and SNOG1del Guide-2 is 3’ to binding site 6 (BS6). Plasmids expressing gRNAs as well as Cas9 and a puromycin resistance cassette were nucleofected into PWS iPSCs. Following transient selection with puromycin, surviving colonies were screened by conventional PCR using primers flanking the intended CRISPR cut sites to identify cells harboring the deletion. Conventional PCR using primers located between the intended cut sites was used to determine whether colonies with the deletion were mixed (i.e. contained both deletion and non-deletion cells) (Supplementary Material, Table S1C). Overall, we identified 2 cell lines carrying a deletion of the entire SNOG1 region in PWS iPSCs. These are termed SNOG1-del1 and SNOG1-del2 (Fig. 1 and Supplementary Material, Table S1B).

Fortunately, the unique sequence flanking the consensus binding motif in each of the six ZNF274 binding sites could be used to specifically target CRISPR/Cas9 to mutate the sites within the *SNORD116* cluster. We designed two different gRNAs to target Cas9 to these specific ZNF274 binding motifs. 116-Z-BS Guide 1 is able to target *SNORD116*-2 to 9 and was expressed transiently using the regular SpCas9 associated with a NGG protospacer adjacent motif (PAM; Fig. 3A, blue box and Supplementary Material, Table S1A). 116-Z-BS Guide 2 was used with the VQR variant of SpCas9 that recognizes a modified PAM sequence NGNG/NGAN. The PAM sequence for this CRISPR encompassed the crucial A-to-G change in the consensus binding motif, which allowed us to target all of the ZNF274 binding sites at the locus without affecting *SNORD116*-2, -4 and -6 (Fig. 3A, red box and Supplementary Material, Table S1A). Following transient delivery of 116-Z-BS Guide 1 and lentiviral delivery of 116-Z-BS Guide 2, puromycin selection was used to eliminate iPSCs that had not received the CRISPR construct. Puromycin resistant colonies were screened via conventional PCR followed by Sanger sequencing for each of the six binding sites (Supplementary Material, Table S1C).

Using the transiently-expressed 116-Z-BS Guide 1 construct, we obtained two cell lines carrying ZNF274 binding site mutations. BS5mut1 harbored a 20 bp deletion within BS5 encompassing 14/18 bp of the ZNF274 consensus binding motif (Fig. 1 and Supplementary Material Fig. S1A and Table S1B). BS6mod-down harbored a 9 bp deletion downstream of the BS6 binding motif (Fig. 1 and Supplementary Material, Fig. S1A and Table S1B). Using the constitutively expressed 116-Z-BS Guide 2 construct, we obtained three cell lines carrying ZNF274 binding site mutations. BS1-4del-BS5mut2 carried a deletion encompassing BS1 to BS4, a 26 bp deletion at BS5 that included 17/18 bp of the ZNF274 consensus binding motif, and a 7 bp insertion upstream of the ZNF274 consensus binding motif in BS6 that only affects the first 2bp of the motif (Fig. 1, Fig. 3A and Supplementary Material, Table S1B). The second cell line, BS5-6mod-up, was found to have a 7 bp deletion at BS5 encompassing the first 5 bp of the ZNF274 consensus binding motif and a 14 bp insertion upstream of the ZNF274 consensus binding motif at BS6 that leaves the entire consensus binding motif intact (Fig. 1 and Supplementary Material Fig. S1A and Table S1B). The third cell line, BS4-5del-BS6mod-up, harbored a deletion spanning BS4 to BS5 and a 6 bp insertion at BS6 that does not affect the ZNF274 consensus binding motif (Fig. 1 and Supplementary Material, Fig. S1A and Table S1B).

### Disruption of *ZNF274* binding sites depletes ZNF274 at the *SNORD116* locus

To determine whether mutating the ZNF274 consensus binding motif affected ZNF274 binding at *SNORD116*, we performed ChIP-qPCR for ZNF274 at BS5, BS6, and a non-*SNORD116* ZNF274 binding locus, *ZNF180* on the PWS iPSC clones carrying various mutations in the ZNF274 binding sites. ChIP-qPCR for these sites were also performed on unedited PWS iPSCs, iPSCs derived from control individuals (CTRL1 and CTRL2)^32; 40-42^, and iPSCs from an AS patient carrying a large deletion of maternal chromosome 15q11-q13^40^ as controls. BS1-4del-BS5mut2 (Fig. 3B), BS5mut1, and BS4-5del-BS6mod-up clones (Supplementary Material, Fig. S1B) showed significantly decreased binding of ZNF274 at BS5, indicating that the BS5 consensus binding motif was severely disrupted or deleted in these clones. Conversely, clone BS5-6mod-up, in which only the first 5 bp of the consensus sequence within BS5 was deleted, showed no significant difference in ZNF274 binding (Supplementary Material, Fig. S1B), indicating that deletion of the first 5 bp is not sufficient to disrupt ZNF274 binding. Using qPCR primers for BS6, there was no significant difference in ZNF274 binding for any of the clones, including clone BS1-4del-BS5mut2, in which the first 2 bp of BS6 were deleted (Supplementary Material, Fig. S1B). For all clones and control iPSCs, binding of the protein at the *ZNF180* 3’UTR was unaffected (Fig. 3B and Supplementary Material, Fig. S1B).

### Disruption of *ZNF274* binding at *SNORD116* restores maternal gene expression in neurons

We first used RT-qPCR to determine whether disruption/deletion of ZNF274 binding sites affected maternal gene expression in PWS iPSCs. We focused on clones carrying deletions of all or most of the ZNF274 consensus motifs. Similar to our previous observations in PWS iPSCs with ZNF274 knocked out ^35^, in BS1-4del-BS5mut2, SNOG1del1 and SNOG2del2 iPSCs, we detected expression using probe-primer sets spanning exons U4 and exon 2 of *SNRPN*, but not exons 1 and 2, suggesting that the alternative upstream promoters but not the canonical promoter of *SNRPN* are activated (Supplementary Material, Fig. S2A). However, this activation of the upstream *SNRPN* exons did not lead to detectable *SNRPN* exon 3/4 or *116HGG2* expression in iPSCs, since the upstream *SNRPN* exons are known to be predominately expressed in neural cell types ^35; 42^.

We next differentiated our engineered PWS iPSCs into neural progenitor cells (NPCs) and forebrain cortical neurons. Consistent with our previous observations quantifying maternal *SNHG14* RNAs in neurons differentiated from ZNF274 knockout iPSCs (LD KO1 and LD KO3), we saw more robust activation of *SNRPN* and *SNORD116* (*SNRPN* ex3/4 and *116HGG2*, respectively) upon neural differentiation of PWS iPSCs with disruptions/deletions in the ZNF274 binding sites (Fig. 4, Supplementary Material, Fig. S2B). In fact, expression levels of these transcripts in NPCs and neurons differentiated from ZNF274 binding site mutated PWS iPSCs was approximately 50% of those seen in NPCs and neurons differentiated from neurotypical iPSCs. Furthermore, NPCs and neurons differentiated from the BS1-4del-BS5mut2 PWS iPSCs, showed equivalent expression levels of these maternal *SNHG14* transcripts as neurons differentiated from SNOG1-del1 and -2 iPSCs. These data further support the hypothesis that ZNF274 binding at maternal *SNORD116* represses neuronal gene expression from the *SNRPN* and *SNHG14*. These data also suggest that that ZNF274 binding to a single site within maternal *SNORD116* is not sufficient to maintain repression of this locus in PWS neurons.

**Figure 4.**
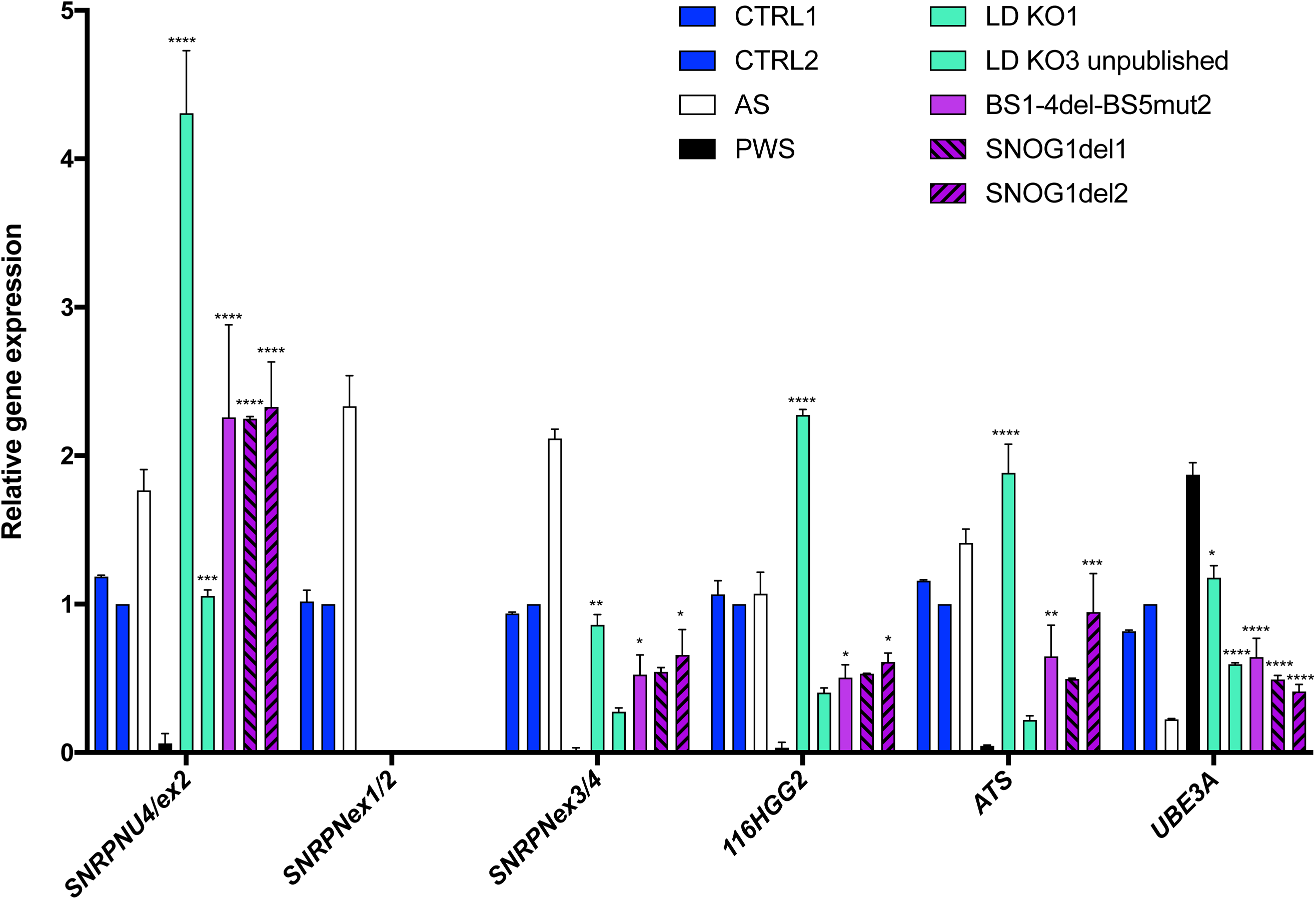
Disrupting ZNF274 binding at *SNORD116* activates transcription in PWS neurons. Expression of the upstream *SNRPN* exons (U4/ex2), *SNRPN* major promoter (ex1/2), *SNRPN* mRNA (ex3/4), the *SNORD116* Host Gene Group II (*116HGG2*), *UBE3A-ATS* and *UBE3A* in iPSCs-derived neurons was quantified using RT-qPCR. Gene expression was assessed using the comparative CT method, *GAPDH* was used as an endogenous control. Data were normalized to CTRL2 for each panel and plotted as the mean with Standard Deviation (SD). A minimum of 2 biological replicates per cell line were performed. Significance was calculated using two-way analysis of variance (ANOVA) test with a Dunnett post-test to compare the disrupted ZNF274 binding cell lines to PWS. *P<0.05, **P<0.01, ***P<0.001, ****P<0.0001.

In NPCs and neurons, expression of the *SNRPN* U4/exon 2 transcripts are fully restored by mutation of the ZNF274 binding sites, while *SNRPN* transcripts that include exon 1 remain silent. Expression levels of the *SNRPN* U4/exon 2 transcripts in PWS NPCs and neurons with mutated ZNF274 binding sites equals or exceeds those seen in neurons differentiated from neurotypical iPSCs, while *SNRPN* exon 3/4 transcripts are only partially activated (Fig. 4, Supplementary Material, Fig. S2B). These results are consistent with our previous work showing that the ZNF274 complex regulates neuronal *SNRPN/SNHG14* transcripts that are initiated from the *SNRPN* upstream promoters.

Disruption of ZNF274 binding also led to expression of *SNHG14* transcripts downstream of *SNORD116* (i.e. *UBE3A-ATS*; Fig. 4) in NPCs and neurons. *UBE3A-ATS* is known to silence paternal *UBE3A* in neurons. Neurons with disrupted ZNF274 binding sites activate *UBE3A-ATS* to ∼50% of normal levels, and *UBE3A* expression is decreased to approximately 50% of normal levels (Fig. 4, Supplementary Material, Fig. S2B). Complete *UBE3A-ATS*-mediated silencing of *UBE3A* may not be observed due to the relative immaturity of the neurons differentiated from the iPSCs. Alternatively, the increased expression of maternal *UBE3A* in PWS iPSC-derived neurons relative to their neurotypical counterparts may counteract the antisense-mediated silencing.

## Discussion

PWS is caused by the loss of paternal gene expression from the chromosome 15q11-q13 locus. Since every individual with PWS has an intact copy of those genes on an epigenetically silenced maternal allele, activating those repressed genes is an attractive therapeutic strategy that addresses the root cause of PWS. The findings summarized here demonstrate that mutation of *ZNF274* consensus binding consensus motifs within maternal *SNORD116* in PWS iPSCs leads to activation of *SNRPN* and *SNHG14* in neurons derived from them. This further supports the notion that prevention of ZNF274 binding at maternal *SNORD116* may be a viable therapeutic approach for PWS.

Identification of the ZNF274 consensus binding motif allowed us to map the precise nucleotides bound by ZNF274 and subsequently design CRISPR constructs to mutate them. Ideally, we would have been able to mutate individual ZNF274 binding sites and identify the minimum number of disrupted sites required to activate *SNHG14* expression. However, our data suggest that binding sites 5 and 6 are the most readily accessible by CRISPR/Cas9, and that deletions of multiple sites along with intervening DNA may be more likely to occur rather than mutating individual internal binding sites (i.e. BS2-4). Sampling a larger number of mutated colonies generated by transiently expressing the 116-Z-BS Guide-1 construct would perhaps have yielded iPSCs harboring more individual binding site mutations. Interestingly, the 116-Z-BS Guide 2 was less efficient at cutting and required constitutive expression via a lentiviral vector to generate mutated ZNF274 binding sites. Although this approach yielded interesting clones, gene expression analyses from neurons differentiated from the more subtle binding site mutations was not possible because these mutations were merely a snapshot in time, and each clone would eventually accumulate more binding site mutations until the gRNA binding was completely abolished from this locus. Similarly, some off-target effects are likely with this approach. Disruption of individual binding sites may be possible with targeted dual CRISPR approaches to flank and delete individual sites one-by-one. Nonetheless, these data strongly suggest that BS5 and BS6 are the most accessible to CRISPR/Cas9.

PWS iPSCs with mutations of BS5 and BS6 allowed us to determine whether ZNF274 binding was disrupted by these mutations. Unsurprisingly, mutations that severely affected the binding sites led to significantly reduced ZNF274 binding, but mutations that removed the first 2-5 bp of the binding site did not significantly affect ZNF274 binding, although ChIP-seq in those iPSCs may provide more accurate quantification of ZNF274 binding in these lines. Interestingly, a G to A nucleotide change at position 8 of the ZNF274 consensus motif that occurs naturally within the human genome is sufficient to prevent ZNF274 binding. These data provide a start to understanding the critical nucleotides in the consensus binding sequence.

Most importantly, by mutating and/or deleting the ZNF274 consensus binding motifs we demonstrated that it is feasible to deplete ZNF274 specifically within *SNORD116* (Fig. 3A,B). The loss of ZNF274 binding at this locus leads to the expression of maternal *SNHG14* in PWS iPSC-derived NPCs and neurons (Fig. 4 and Supplementary Material, Fig. S2A,B). The expression levels of these activated transcripts approach normal levels and robust activation is observed not only observed within the *SNORD116* portion of *SNHG14*, but also extends throughout the proximal and distal portions of the *SNHG14* RNA, as shown by *SNRPN* and *UBE3A-ATS* expression (Fig. 4 and Supplementary Material, Fig. S2A,B).

The canonical promoter of *SNRPN* was not activated by *ZNF274* binding disruption (Fig. 4 and Supplementary Material, Fig. S2A,B). This was previously observed in PWS iPSCs carrying a full knockout of *ZNF274*, as well. We previously demonstrated that these ZNF274 knockout iPSCs did not have altered CpG methylation at the maternal PWS-IC compared to unedited PWS iPSCs. These data show that removal of ZNF274 binding at *SNORD116* does not affect DNA methylation at the PWS-IC and does not activate the canonical *SNRPN* promoter^35^. Instead, disruption of ZNF274 binding at *SNORD116* leads to activation of upstream *SNRPN* promoters. These promoters are preferentially expressed in NPCs and neurons. We observe expression levels of upstream *SNRPN* transcripts in ZNF274 binding site-mutated PWS NPCs and neurons that are similar to or even exceed those seen in neurotypical NPCs and neurons. These data further support the hypothesis that ZNF274 binding to maternal *SNORD116* serves as a somatic imprint to maintain repression of *SNRPN* and *SNHG14* in neural lineages.

As previously observed with our *ZNF274* knockout PWS iPSCs, we did not detect substantially decreased levels of *UBE3A* despite activation of *UBE3A-ATS* (Fig. 4 and Supplementary Material, Fig. S2A,B). It is possible that *UBE3A-ATS*-mediated silencing of *UBE3A* may not be detectable due to the relative immaturity of the neurons differentiated from the iPSCs compared to a fully developed brain.^40^ Alternatively, it is possible that the levels of expression of the maternal *UBE3A* mRNA are balanced with those of the *UBE3A-ATS* transcript and thus we do not see full antisense-mediated silencing.

While it is clear that ZNF274 plays an important role in mediating the repression of the upstream *SNRPN* promoters in neurons, the specific histone methyltransferases and other co-factors involved are not as certain. We previously implicated the H3K9me3 histone methyltransferase, SETDB1, in this process and showed that PWS iPSCs with a knockdown of SETDB1 also activated maternal *SNHG14* and *SNRPN* ^32^. SETDB1 is a well-known partner of ZNF274 ^33^. Interestingly, Kim et al successfully activated maternal *SNRPN* and *SNHG14* in human PWS fibroblasts and a mouse model of PWS, using novel compounds that inhibit the histone methyltransferase G9a ^43; 44^. This activation of maternal PWS RNAs via G9a inhibition was linked to reduced levels of H3K9me3 and H3K9me2 at the *SNORD116* locus as well as reduced levels of H3K9me2 at the PWS-IC, without affecting DNA methylation levels at the PWS-IC ^43^. Similarly Wu et al. showed activation of *SNHG14* and *SNRPN* in human PWS iPSC-derived NPCs and neurons using G9a inhibitors (https://www.biorxiv.org/content/10.1101/640938v1). Although the association of G9a with ZNF274 has not previously been shown, G9a and SETDB1 have been reported to complex together ^45^. Whether the G9a- and the ZNF274/SETDB1 complex-mediated H3K9me3 silencing of maternal chromosome 15q11-q13 transcripts are redundant or complimentary remains unknown. It will be important to determine the number of other genes affected by SETDB1, G9a, and ZNF274 individually, and the extent to which the targets of these epigenetic regulators interact both to better understand the repressive mechanisms working on the *SNORD116* locus, but also to identify the potential pitfalls of SETDB1, G9a, or ZNF274 inhibition as therapeutic approaches for PWS, such as affecting non-PWS related genes ^36; 46^. Fortunately, our results show the feasibility of disrupting ZNF274 binding specifically at the maternal *SNORD116* locus. We hypothesize that this targeted approach will lead to restoration of appropriate *SNRPN/SNHG14* gene expression without impacting other genes, providing a safer approach compared to inhibition of major epigenetic regulators. Further investigation into how to best prevent ZNF274 from binding at maternal *SNORD116* is needed to better define a potential strategy for future therapeutic application for PWS.

## Material and Methods

### Culture conditions of iPSCs and neuronal differentiation

iPSCs were grown on irradiated mouse embryonic fibroblasts and fed daily with conventional hESC medium composed of DMEM-F12 supplemented with knock-out serum replacer, nonessential amino acids, L-glutamine, β-mercaptoethanol, and basic FGF. iPSCs were cultured in a humid incubator at 37°C with 5% CO_2_ and manually passaged once a week ^40^.

Neuronal differentiation of iPSCs was performed using a monolayer differentiation protocol ^47; 48^ with some modifications ^40; 41^. Briefly, iPSC colonies were cultured in hESC medium for 24h before switching to N2B27 medium. Cells were fed every other day with N2B27 medium containing Neurobasal Medium, 2% B-27 supplement, 2mM L-glutamine, 1% Insulin-transferrin-selenium, 1% N2 supplement, 0.5% Pen-strep and was supplemented with fresh noggin at 500ng/mL. After three weeks of neural differentiation, neural progenitors were plated on tissue culture plates coated with poly-ornithine/laminin. The neural differentiation medium consisted of Neurobasal Medium, B-27 supplement, nonessential amino acids, and L-glutamine, and was supplemented with 1 μM ascorbic acid, 200 μM cyclic adenosine monophosphate, 10 ng/mL brain-derived neurotrophic factor, and 10 ng/mL glial-derived neurotrophic factor. Unless otherwise specified, cells were harvested once neural cultures reached at least 10 weeks of age.

### Lentiviral production, transduction, and clone screening

sgRNAs were designed using a web-based CRISPR design tool and cloned into lentiCRISPR (Addgene Plasmid 49535 and 52961) original or modified to create the VQR mutation, lentiGuidePuro (Addgene Plasmid 52963) or pX459 v2.0 (Addgene plasmid 62988) using our standard protocol ^49-51^. Lentiviral particles were made by transfecting 293FT cells with 2^nd^ generation packaging systems using Lipofectamine 2000 (Life Technologies). Prior to transduction or electroporation, iPSCs were treated with 10 μM ROCK inhibitor, Y-27632, overnight. The next day, iPSCs were singlized using Accutase (Millipore) before transduction/electroporation. Transduction was done with lentivirus in suspension in the presence of 8 μg/mL polybrene in a low-attachment dish for two hours. Then, the iPSCs/lentivirus mixture was diluted 1:1 in hESC medium before plating. Electroporation was performed in 0.4cm cuvettes loaded with 10µg of the CRISPR/Cas9 and 800µL of PBS suspended iPSCs. Cells were electroporated using a Biorad Gene Pulser X Cell with the exponential protocol, at 250V, a 500µF capacitance, ∞ resistance. Transduced/electroporated cells were plated on puromycin-resistant (DR4) MEF feeders at a low density, supplemented with 10 μM ROCK inhibitor, Y-27632, overnight. Following transduction, attached cells were cultured in hESC medium for an additional 72 hours before starting drug selection using puromycin at 0.5 μg/mL during the first week and at 1 μg/mL during the second week. Following electroporation, at 24 hours post plating, the cells were selected with 0.5 μg/mL of puromycin for a total of 48 hours. Puromycin-resistant iPSC colonies were individually picked into a new feeder well and screened for indels by performing PCR on genomic DNA and sequencing. The sgRNA sequences and PAM are summarized in Supplementary Material, Table S1A. The genetic alterations induced are detailed in Fig. 1, Fig. 3A and Supplementary Material, Fig. S1A. The cell lines are summarized in Supplementary Material, Table S1B. PCR primers used to amplify the desired genomic regions are summarized in Supplementary Material, Table S1C.

### RNA isolation and RT reaction

RNA was isolated from cells using RNA-Bee (Tel Test, Inc.). Samples were DNase-treated as needed with Amplification Grade DNaseI (Invitrogen) at 37°C for 45 minutes, and cDNA was synthesized using the High Capacity cDNA Reverse Transcription Kit (Life Technologies) according to the manufacturer’s instructions.

### RT-qPCR and expression arrays

For single gene expression assays, expression levels of target genes were examined using TaqMan Gene Expression Assays (Applied Biosystems) on the Step One Plus (ThermoFisher Scientific) or on the BioRAD CFX96 Real Time PCR system (Biorad). An amount of RT reaction corresponding to 30ng of RNA was used in a volume of 20ul per reaction. Reactions were performed in technical duplicates or triplicates and the *GAPDH* Endogenous Control TaqMan Assay was used as an endogenous control, following the manufacturer’s protocol. Relative quantity (RQ) value was calculated as 2^− ΔΔCt^ using the normal cell lines CTRL1 or CTRL2 as the calibrator sample.

### Chromatin Immunoprecipitation (ChIP)

ChIP assays were performed as described before ^32; 35; 52; 53^. The antibody anti-ZNF274 (Abnova, Cat# H00010782-M01) was used. Quantification of ChIPs was performed using SYBR Green quantitative PCR. PCR primers used to amplify the purified DNA can be found in Supplementary Material, Table S1C. The enrichment of the DNA was calculated as percent input, as described.^53^ Normal rabbit IgG was used for the isotype controls and showed no enrichment. Data were presented as means with SD and represent the average of at least two biological replicates from independent cultures.

### Statistical tests

Statistical analysis was carried out using Prism software (GraphPad). For each condition shown, averaged values from a minimum of two biological replicates from independent cultures were calculated and the resulting standard deviation (SD) was reported in the error bars. Unless otherwise specified, for each experiment, averaged values for each sample were compared to that of the parental PWS cell line of the same genotype (PWS LD) and the significance for each un-manipulated vs. KO pair was calculated using the one- or two-way analysis of variance (ANOVA) with the Dunnett post-test.

## Acknowledgments

We thank David S. Rosenblatt, Gail Dunbar and Daniel J Driscoll for patient clinical evaluation and information, and for providing skin biopsies/fibroblasts. We thank the UCONN Health Molecular Core. We thank James A. Thomson, John P. Maufort, Elizabeth S. Perrin, and Jessica Antosiewicz-Bourget at Wisconsin National Primate Research Center, University of Wisconsin– Madison for the cynomolgus iPSCs and for technical assistance with cynomolgus iPSCs culture. This work was supported by the Foundation for Prader-Willi Research and the CT Regenerative Medicine Fund (to M. Lalande), the Cascade fellowship (to M. Langouët) and Levo Therapeutics (to S. Chamberlain). The contents in this work are solely the responsibility of the authors and do not necessarily represent the official views of the state of Connecticut.

## Supplemental Data

Supplemental Data include 2 figures and 3 tables and can be found with this article online.

## Web Resources

UCSC Human Genome Browser, http://genome.ucsc.edu/cgi-bin/hgGateway

Web-based CRISPR design tool, http://crispr.mit.edu

TIDE: method for easy quantitative assessment of genome editing, https://tide.nki.nl/

CRISP-ID: Detecting CRISPR mediated indels by Sanger sequencing, http://crispid.gbiomed.kuleuven.be/

RoadMap Epigenomics, http://egg2.wustl.edu/roadmap/web_portal/imputed.html#imp_sig

## Author Contributions

Maéva Langouët (M.L.) and J.C. analyzed the ChIP-seq data and J.C. identified the consensus binding motif for ZNF274. M.L., C.O., C.D.T., H.G.D. and C.S. designed and tested the CRISPR/gRNAs. M.L., C.O. and D.G. screened and generated the engineered cell lines. M.L., C.O., D.G., M.C. and L.C. characterized the engineered cell lines. M.L. executed and analyzed ChIP data from human iPSCs. M.C. executed and analyzed ChIP data from Cynomolgous stem cells. M.L., N.G. and D.G. performed neuronal differentiation. M.L. and D.G. performed and analyzed the gene expression assays. M.L. executed statistical analysis of the data. M.L., C.S., S.C. and M.Lalande designed and directed the study. All authors contributed to writing and editing the manuscript.

## Supplementary Information

**Figure S1. ZNF274 binding in engineered PWS iPSCs**.

**A**. Sequences of group 1 *SNORD116* copies are shown. The ZNF274 consensus sequence identified here is highlighted in yellow. The position of the ZNF274 motif proposed by Imbeault et al. is indicated. *SNORD116* copies bound by ZNF274 are in black font, while those not bound by ZNF274 are in gray font. Single base substitutions are highlighted in colored fonts. and the corresponding ZNF274 ChIP assays **B**. ZNF274 ChIP assays for iPSCs in A. Quantification of ChIP was performed and calculated as percent input for each sample. Binding at *ZNF180* is included as a positive control. Samples were normalized against the PWS (black) sample. A minimum of 2 biological replicates per cell line were performed. Significance was calculated using two-way analysis of variance (ANOVA) test with a Dunnett post-test to compare the disrupted ZNF274 binding cell lines to PWS. *P<0.05, **P<0.01.

**Figure S2. Levels of maternal genes activation in PWS iPSCs and NPCs following disruption of ZNF274 binding at *SNORD116***.

**A**. Expression of the upstream *SNRPN* exons (U4/ex2), *SNRPN* major promoter (ex1/2), *SNRPN* mRNA (ex3/4), the *SNORD116* Host Gene Group II (*116HGG2*) and *UBE3A* in iPSCs and **B**. and NPCs.

## Supplemental Tables

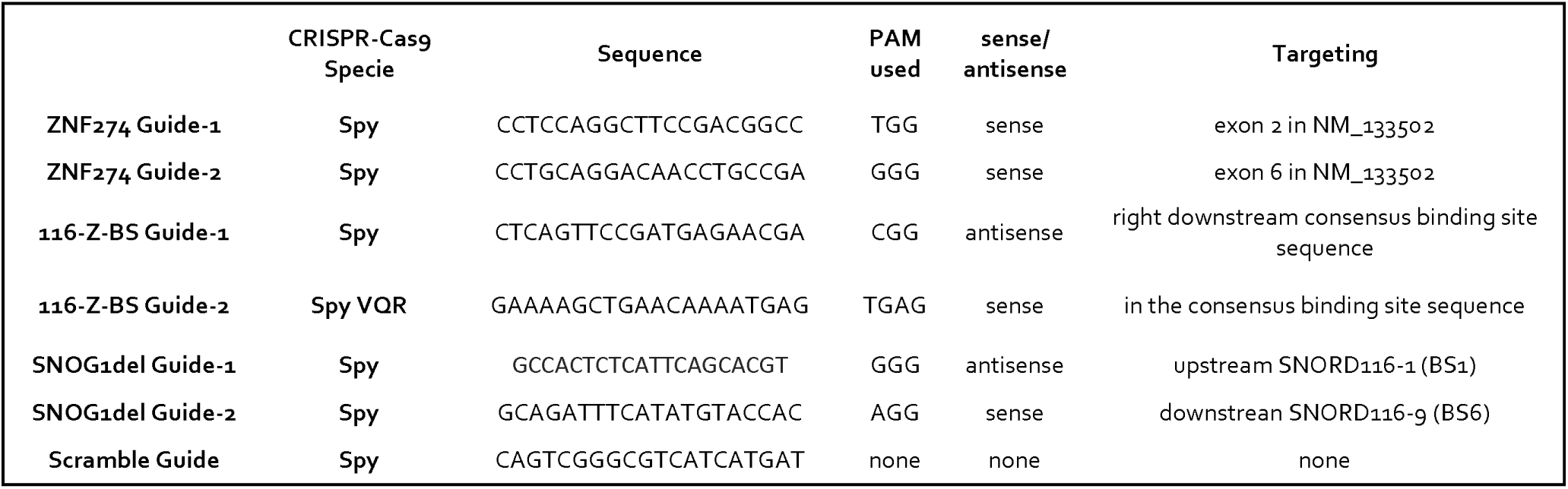
A.

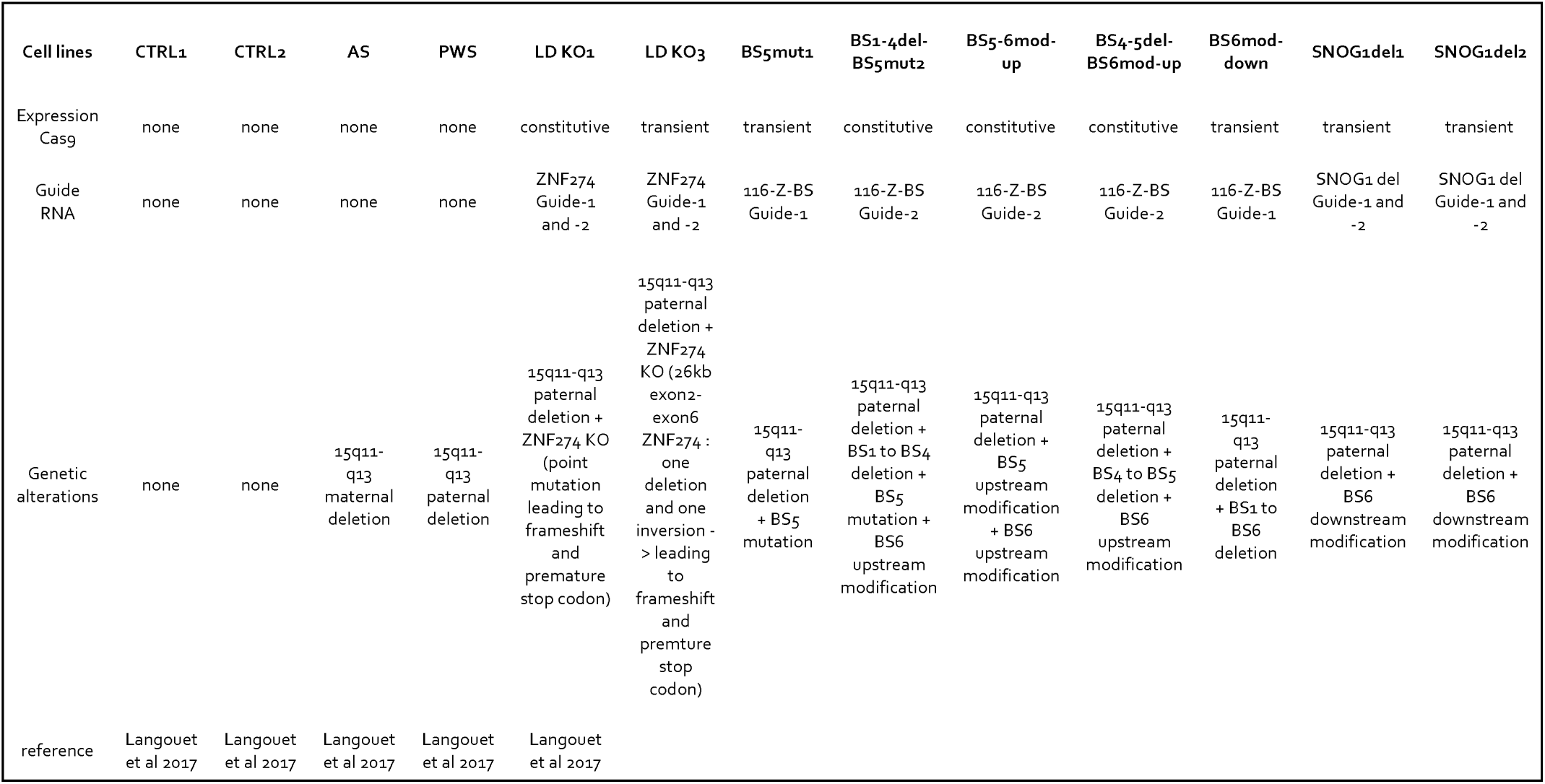
B.

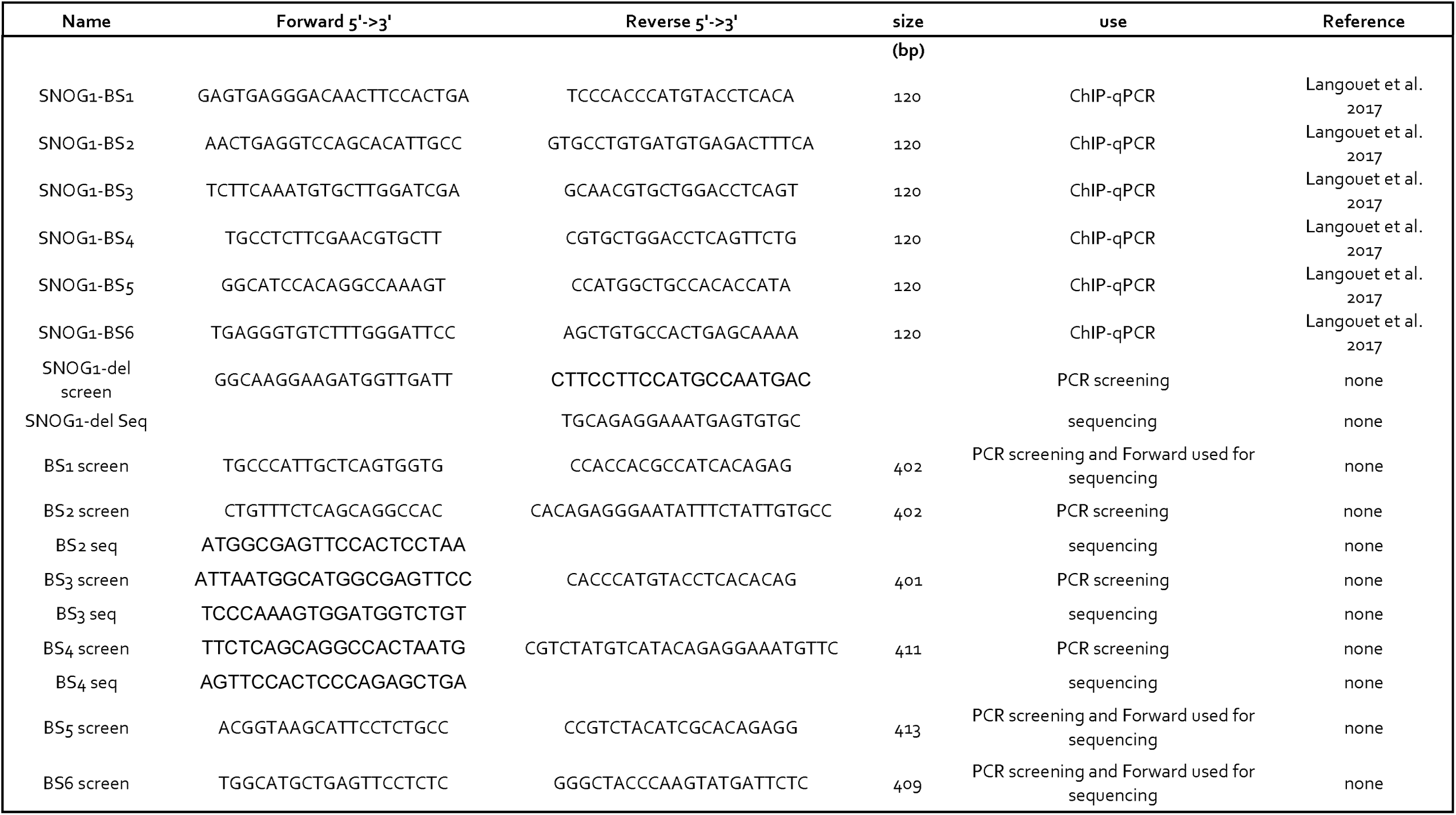
C.

## Abbreviations

116HGG2: SNORD116 host gene Group2 transcript
3’UTR: 3’ Untranslated Transcribed Region
AS: Angelman syndrome
ChIP: Chromatin ImmunoPrecipitation
CRISPR: Clustered Regularly Interspaced Short Palindromic Repeats
Cas9: CRISPR associated protein 9
CTRL: iPSCs from control individuals
G9a: histone methyltransferase
H3K9me2: histone H3 lysine 9 dimethylation
H3K9me3: histone H3 lysine 9 trimethylation
HG: host gene
iPSCs: induced pluripotent stem cells
lncRNA: long non-coding RNA
NPCs: neural progenitor cells
PWS: Prader-Willi syndrome
PWS-IC: PWS-Imprinting Center
SETDB1: SET domain bifurcated 1
SNOG1: SNORD116 Group 1
SNOG2: SNORD116 Group 2
SNOG3: SNORD116 Group 3
SNORD115: box C/D class small nucleolar RNAs
SNORD116: box C/D class small nucleolar RNAs
SNRPN: small nuclear ribonucleoprotein polypeptide N
UBE3A: Ubiquitin Protein Ligase E3A
UBE3A-ATS: antisense overlapping UBE3A transcript
ZNF274: zinc-finger protein ZNF274
ZNF274 BS: ZNF274 binding sites
LD KO1 & 3: ZNF274 knockout from PWS large deletion (LD) iPSCs

