## Supplementary figures and images for "Specific ZNF274 binding interference at *SNORD116* activates the maternal transcripts in Prader-Willi syndrome neurons"

### Supplemental FigS2

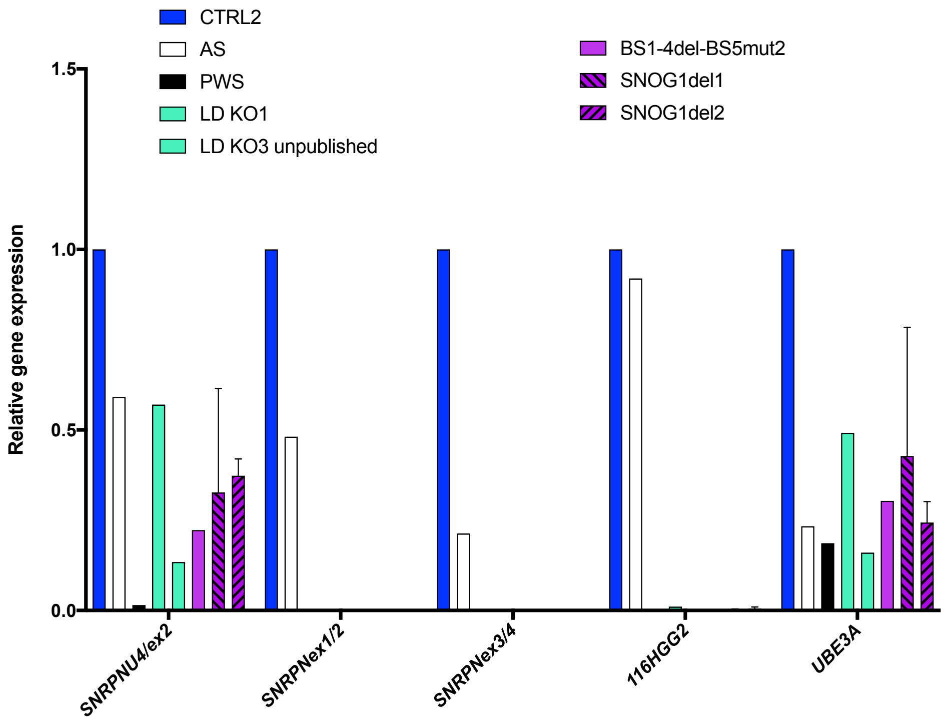

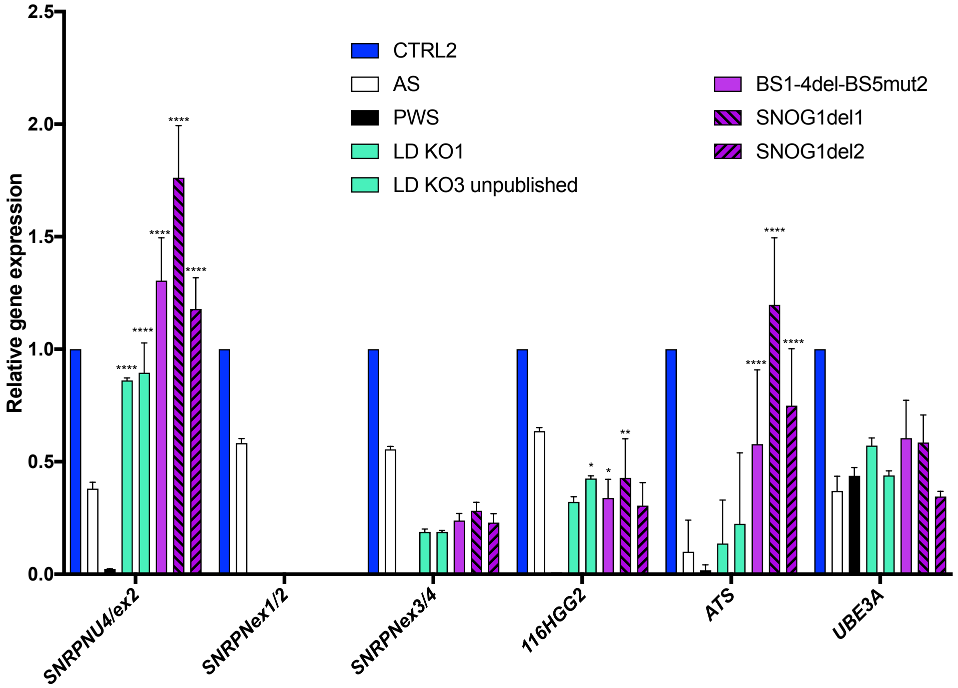
