## Supplemental FigS1 for "Specific ZNF274 binding interference at *SNORD116* activates the maternal transcripts in Prader-Willi syndrome neurons"

## A

### BS5mut1 (116-Z-BS Guide-1)

**BS1** *SNORD116-1* ...AAAAACATTCCTTGGAAAAGCTGAACAAAATGAGTGAGAACTCATAACGTCATTCTCATCGGAACTGAGGTCCAGCA...

*SNORD116-2* ... AAAAACATTCCTTGGAAAAGCTGAACAAAATGAGTGAAAACTCATACCGTCATTCTCATCGGAACTGAGGTCCAGCA...

**BS2** *SNORD116-3* ...AAAAACATTCCTTGGAAAAGCTGAACAAAATGAGTGAGAACTCATACCGTCGTTCTCATCGGAACTGAGGTCCAGCA...

*SNORD116-4* ...AAAAACATTCCTTGGAAAAGCTGAACAAAATGAGTGAAAACTCATACCGTCGTTCTCATCGGAACTGAGGTCCAGCA...

**BS3** *SNORD116-5* ...AAAAACATTCCTTGGAAAAGCTGAACAAAATGAGTGAGAACTCATACCGTCGTTCTCATCAGAACTGAGGTCCAGCA...

*SNORD116-6* ...AAAAACATTCCTTGGAAAAGCTGAACAAAATGAGTGAAAACTCATACCGTCATTCTCATCGGAACTGAGGTCCAGCA...

**BS4** *SNORD116-7* ...AAAAACATTCCTTGGAAAAGCTGAACAAAATGAGTGAGAACTCATACCGTCGTTCTCATCAGAACTGAGGTCCAGCA...

**BS5** ***SNORD116-8*** ...AAAAACATTCCTTGGAAAAGCTGAACAAAATGAG*********************CTCATCGGAACTGAGGTCCAGCA...

**BS6** *SNORD116-9* ...AAAAACATTCCTTGGAAAAGCTGAACAAAATGAGTGAGAACTCATACCGTCGTTCTCATCGGAACTGAGGTCCAGCA...

### BS6mod-down (116-Z-BS Guide-1)

**BS1** *SNORD116-1* ...AAAAACATTCCTTGGAAAAGCTGAACAAAATGAGTGAGAACTCATAACGTCATTCTCATCGGAACTGAGGTCCAGCA...

*SNORD116-2* ... AAAAACATTCCTTGGAAAAGCTGAACAAAATGAGTGAAAACTCATACCGTCATTCTCATCGGAACTGAGGTCCAGCA...

**BS2** *SNORD116-3* ...AAAAACATTCCTTGGAAAAGCTGAACAAAATGAGTGAGAACTCATACCGTCGTTCTCATCGGAACTGAGGTCCAGCA...

*SNORD116-4* ...AAAAACATTCCTTGGAAAAGCTGAACAAAATGAGTGAAAACTCATACCGTCGTTCTCATCGGAACTGAGGTCCAGCA...

**BS3** *SNORD116-5* ...AAAAACATTCCTTGGAAAAGCTGAACAAAATGAGTGAGAACTCATACCGTCGTTCTCATCAGAACTGAGGTCCAGCA...

*SNORD116-6* ...AAAAACATTCCTTGGAAAAGCTGAACAAAATGAGTGAAAACTCATACCGTCATTCTCATCGGAACTGAGGTCCAGCA...

**BS4** *SNORD116-7* ...AAAAACATTCCTTGGAAAAGCTGAACAAAATGAGTGAGAACTCATACCGTCGTTCTCATCAGAACTGAGGTCCAGCA...

**BS5** *SNORD116-8* ...AAAAACATTCCTTGGAAAAGCTGAACAAAATGAGTGAGAACTCATACCGTCGTTCTCATCGGAACTGAGGTCCAGCA...

**BS6** ***SNORD116-9*** ...AAAAACATTCCTTGGAAAAGCTGAACAAAATGAGTGAGAACTCATACCGTCG*********GAACTGAGGTCCAGCA...

### BS5-6mod-up (116-Z-BS Guide-2)

**BS1** *SNORD116-1* ...AAAAACATTCCTTGGAAAAGCTGAACAAAA-----------------------TGAGTGAGAACTCATAACGTCATTCTCATCG

*SNORD116-2* ... AAAAACATTCCTTGGAAAAGCTGAACAAAA----------------------TGAGTGAAAACTCATACCGTCATTCTCATCG

**BS2** *SNORD116-3* ...AAAAACATTCCTTGGAAAAGCTGAACAAAA----------------------TGAGTGAGAACTCATACCGTCGTTCTCATCG

*SNORD116-4* ...AAAAACATTCCTTGGAAAAGCTGAACAAAA-----------------------TGAGTGAAAACTCATACCGTCGTTCTCATCG

**BS3** *SNORD116-5* ...AAAAACATTCCTTGGAAAAGCTGAACAAAA----------------------TGAGTGAGAACTCATACCGTCGTTCTCATCA

*SNORD116-6* ...AAAAACATTCCTTGGAAAAGCTGAACAAAA----------------------TGAGTGAAAACTCATACCGTCATTCTCATCG

**BS4** *SNORD116-7* ...AAAAACATTCCTTGGAAAAGCTGAACAAAA-----------------------TGAGTGAGAACTCATACCGTCGTTCTCATCA

**BS5** ***SNORD116-8*** ...AAAAACATTCCTTGGAAAAGCTGAACAA**-----------------------******GAGAACTCATACCGTCGTTCTCATCG

**BS6** ***SNORD116-9*** ...AAAAACATTCCTTGGAAAAGCTGAACAAAACTTTTGTTTTGTTATGAGTGAGAACTCATACCGTCGTTCTCATCG

### BS4-5del-BS6mod-up (116-Z-BS Guide-2)

**BS1** *SNORD116-1* ...AAAAACATTCCTTGGAAAAGCTGAACAAAA-----------TGAGTGAGAACTCATAACGTCATTCTCATCGTTCTCATCGGAACTG

*SNORD116-2* ... AAAAACATTCCTTGGAAAAGCTGAACAAAA-----------TGAGTGAAAACTCATACCGTCATTCTCATCGTTCTCATCGGAACTG

**BS2** *SNORD116-3* ...AAAAACATTCCTTGGAAAAGCTGAACAAAA-----------TGAGTGAGAACTCATACCGTCGTTCTCATCGGAACTGAGGTCCAG

*SNORD116-4* ...AAAAACATTCCTTGGAAAAGCTGAACAAAA------------TGAGTGAAAACTCATACCGTCGTTCTCATCGGAACTGAGGTCCAGC

**BS3** *SNORD116-5* ...AAAAACATTCCTTGGAAAAGCTGAACAAAA-----------TGAGTGAGAACTCATACCGTCGTTCTCATCAGAACTGAGGTCCAG

*SNORD116-6* ...AAAAACATTCCTTGGAAAAGCTGAACAAAA------------TGAGTGAAAACTCATACCGTCATTCTCATCGGAACTGAGGTCCAGC

**BS4** *SNORD116-7*

**BS5** *SNORD116-8*

**BS6** ***SNORD116-9*** ...AAAAACATTCCTTGGAAAAGCTGAACAAAACCGGTATGAGTGAGAACTCATACCGTCGTTCTCATCGGAACTGAGGTCCAG

**B**

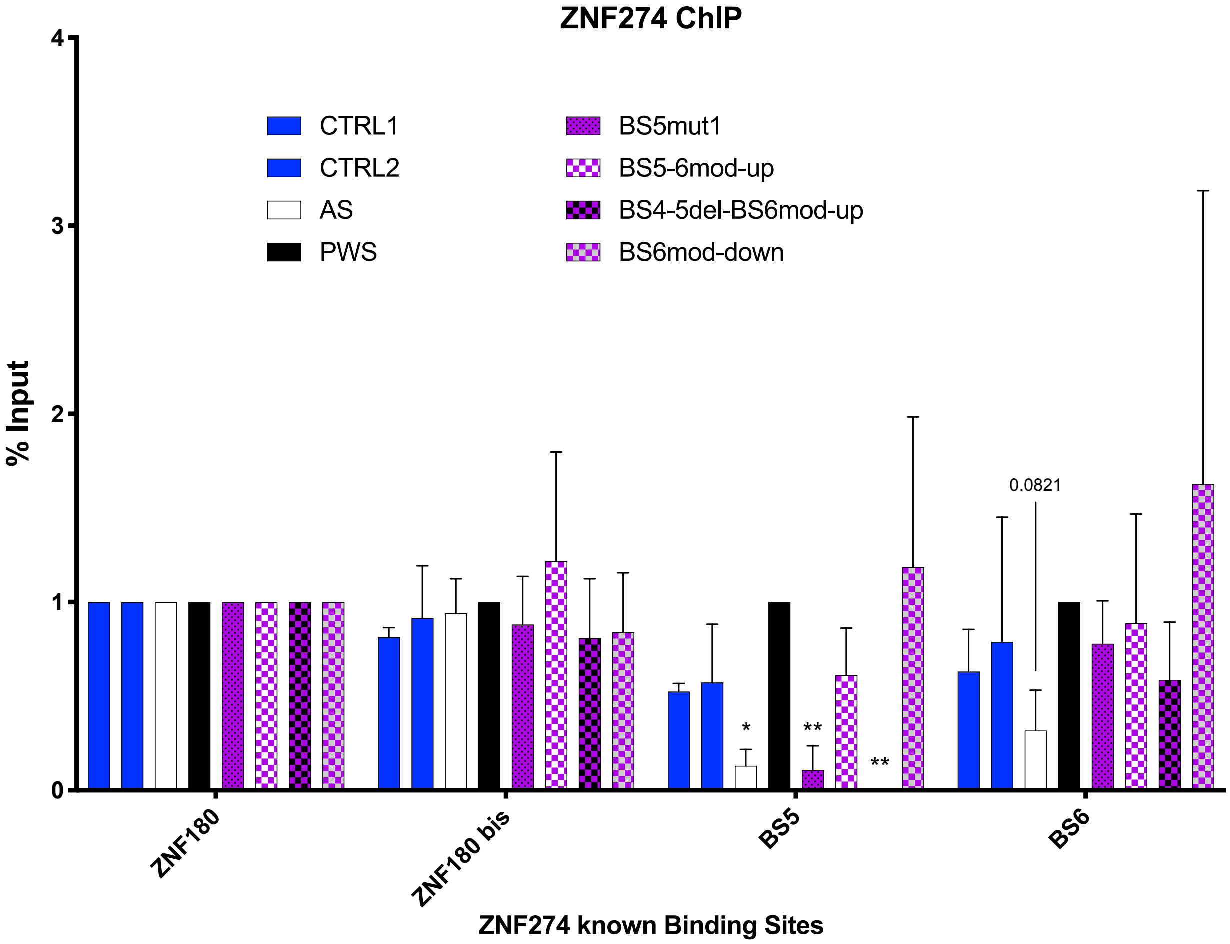
ZNF274 ChIP
% Input
0
1
2
3
4
CTRL1
CTRL2
AS
PWS
BS5mut1
BS5-6mod-up
BS4-5del-BS6mod-up
BS6mod-down
*
**
**
0.0821
ZNF180
ZNF180 bis
BS5
BS6
ZNF274 known Binding Sites
